# The different functions of exercise, supplementary resveratrol, and the combination of exercise and supplementary resveratrol in young and old SAMP8 mice liver

**DOI:** 10.1101/2020.07.19.210542

**Authors:** Sergey Suchkov, Jia-Ping Wu

## Abstract

The senescence-accelerated mouse prone 8 (SAMP8) has about half the normal lifespan of a rodent. Therefore, we determined whether the different age degrees has rapid physiological senescence with the same interaction function and mechanism of exercise and supplementary resveratrol intake by SAMP8 mice liver. Histological of aging related liver disease was examined SAMP8 mice by hematoxylin-eosin and Masson’s trichrome staining.

Apoptosis was determined by TUNEL staining. The mechanism proteins expression of p-PI3K/PI3K, p-Akt/Akt, Bad, cytochrome c, Bcl 2, ERK1, IL-6, STAT3, MEK5, p-ERK5/ERK5, FGF2, and MMP2/9 were examined by western blot. Results showed that the 3- and 6-month-old SAMP8 mice liver, the senescence-accelerated liver cross-sections observed adipocytes, collagen, and apoptosis cells appear, but lower occurs by exercise, supplementary resveratrol, and their combination. Combination exercise and supplementary resveratrol at the 6-month-old SAMP8 mice has significant increase ratio by p-PI3K/PI3K compared to 3-month-old SAMP8 mice liver (p<0.01), and compared with without treatment SAMP8 mice liver. Exercise, supplementary resveratrol intake, and combined of exercise and resveratrol has facilitated p-Akt/Akt ratio increases, and ERK1/Bcl2 increases, but Bad/Cytochrome c decreases at the 6-month-old SAMP8 by western blot. IL-6/STAT3/MEK5 and p-ERK5/ERK5 ratio have good functions at 6-month-old SAMP8 mice better than 3-month-old SAMP8. FGF2/MMP2 decreases and MMP9 increases observed at the 3-month-old SAMP8 by exercise, supplementary resveratrol, and combination. Indeed, we suggest supplementary resveratrol can help exercise therapy ageing related liver diseases.

## Introduction

Aging can divide into two categories: genetic programs and stochastic environment risks. The aging is an age-dependent physiological from the born and raised in (Chen et al., 2019). Senescence-accelerated mice (SAM) has showed observation features indicative of accelerated aging such as hair loss, loss of activity, lordokyphosis, and shortened life (Takigawa et al., 2019). However, littermates of the SAMP mice did not show senescence-related phenotypes were bred, the main characteristic of young SAMP mice keep normal development and waited to maturity of reproductive function, the old SAMP mice has an early manifestation of senescence-related phenotypes. Pathological phenotypes observed in old SAMP8 are associated. The severity diseases increases with aging (Yuan et al., 2018). Therefore, we want to know whether age-related disorder in SAMP8 mice is a physiological senescence rather than pathological. Do you want to live a longer life in good health? Simple practices can make some difference, such as exercise or nutrient supplementary. The liver is essential in the whole-body metabolism function through alterations of multiple metabolic pathways. Both Resveratrol (Re, 3,5,4’-trihydroxy-trans-stilbene) and exercise maybe reverse the effect of aging in the liver maintained high activities of higher antioxidant and decreasing oxidative damage (Adam et al., 2013). Resveratrol (Re) is a stilbenoid compound produce, a type of natural phenol, with many biologic activities. The antiaging effects of resveratrol on several biologic actions such as modulate different metabolic signaling pathways. Resveratrol also shows protective activity against age related deterioration in SAMP8 mice (Adamson, 2013). Exercise training has been shown to have multiple beneficial effects on metabolism in the liver. Regular exercise maintains human physiological function. Exercise rehabilitation regimen helps physiologic and pharmacologic changes is important in returns to normal on organ functions, including reversed aging (Ambrose and Barua, 2004). We examined the effects of exercise, resveratrol, and combination exercise and supplementary resveratrol on the aging associated decline in SAMP8 mice. However, whether lifelong exercise training combination with resveratrol can prevent the age-associated change still remains to be elucidated. As a biological individual, this process has the different molecular mechanisms to regulate different aging grade. The effects of resveratrol on exercise are required for further investigation. Resveratrol intake plays a critical pathophysiological role in maintaining and subverting liver function in the setting of progressive fibrosis.

## Materials and Methods

### SAMP8 mice animal model

It is suggested that purchased 3-month-old male SAMP8 mice should be randomly divided into a young control group and 3 subjects’ groups (exercise, resveratrol intake, combined exercise and resveratrol). The 6-month-old SAMP8 mice as mature old mice. Beginning to test experiments, animals need to keep more than 12 weeks, and then compared with the control group of aging biological activity indicators. Male SAMP8 mice were purchased from BioLASCO TAIWAN at 6 weeks of age and then fed to 3-month-old and 6-month-old male mice at the Animal Experimental Center of China Medical University. The use of the Taiwan Council approved the animal care and experiment. The animal experimental guidelines use of Taipei Medical University Animal Care and Use Committee (LAC-2019-0264) were followed. All mice are housed in a cage in an environment controlled animal room. The temperature of the animal compartment was maintained at 25 °C, the relative humidity was about 55~60%.

### Animals Study designs

The study design was examined by aging mice exercise training or received chronic resveratrol treatment on aging-related diseases. SAMP8 mice promoted aged 3 months (young) and 6 months (old) treated exercise, supplementary resveratrol intake, and combined of exercise and resveratrol to anti-aging. Exercise training consisted of running for 30 min/day, 3 days, one week at a temperature of 25 °C for 4 weeks. The 3-month-old (young) and 6-month-old (old) SAMP8 mice were supplementary resveratrol intake treatment 20 mg/kg, one day, 4 weeks. Animals were fed twice daily for four weeks. Carried out this experimental, resveratrol intake (20 mg/kg/day, oral dosing) were the period of four weeks during running roller wheel introduced to 3-month-old (young) and 6-month-old (old) SAMP8 mice. All animals were sacrificed after the last exercise session.

### Hematoxylin-eosin, Masson’s trichrome, and TUNEL staining

Liver sections were deparaffinized. They are from 100% of a series of graded alcohols to 90% to 70% for 5 minutes each time after and then treated to hematoxylin-eosin (H&E), Masson’s trichrome, TUNEL assay (Apoptosis detection kit, TA300, R&D Systems, Minneapolis, MN, USA) following supplier instructions. After washing with phosphate-buffered saline (PBS), each slide was then soaked with 75%, 85%, 95%, 100% alcohol. Cross-section of the color image with the Nikon E600 in the total magnification of 200x and 40x. Microscope lighting is optimized by visualization in the appropriate cross-section. Determination of the cross-sectional area of collagen and apoptosis cells in the amount and evaluation of statistical analysis.

### Western blotting analysis

Each liver sample (50 μg/lane) was separated via 12.5% SDS-polyacrylamide gel electrophoresis conducted at 110□V voltage for 80□min, 1X Tris–Glycine-SDS running buffer then transferred onto PVDF membranes (Bio-Rad, CA, USA) for 2□h at 400□mA constant 1X Tris–Glycine buffer containing 20% methanol. After transfer PVDF membranes were washed with tris-buffered saline (TBS) and then blocked with 5% bovine serum albumin (BSA, Sigma-Aldrich, Missouri, USA) in 1X TBS with 0.1% Tween-20 (TBS-T), 1□h at room temperature. Incubated with primary mouse antibody against anti-p-PI3K mAb, anti-PI3K mAb, anti-p-Akt mAb, anti-Akt mAb, anti-Bcl2 mAb, anti-Bad mAb, anti-cytochrome c mAb, 1:550, (BD Biosciences, San Jose, CA, USA); anti-ERK1 mAb, anti-STAT3 mAb, anti-MEK5 mAb, 1:250, (Cell Signaling, Danvers, MA, USA); anti-IL-6 mAb, anti-p-ERK5 mAb, anti-ERK5 mAb, anti-tubulin mAb, 1:500, (Promega, Madison, Wisconsin) overnight at 4°C. After washing, the membranes were incubated with an HRP-conjugated secondary antibody, anti-mouse IgG, 1:1000, (Invitrogen, Waltham, Massachusetts, USA) and enhanced chemiluminescence (Amersham Pharmacia Biotech, Piscataway, NJ) was applied for 5□min. Blots were imaged using a single 60□s exposure on a Kodak camera (Gel Logic 1500 Imaging System) and bands band density was performed using Image J program.

### Statistical Analysis

Counting by scanning using FUJIFILM Imagine the intensity of the hybridization signal in the Western blot analysis program. Statistical data was performed using SigmaStat10 software. All results are expressed as mean ± SEM. Statistical analysis was performed using one-way ANOVA following student t-test analysis. When evaluating multiple groups, two-way ANOVA was used with student t-test. Statistical analysis was determined by student’s unpaired t-test (two-tailed), when the 6-month-old old genetic type of SAMP8 mice liver compared with 3-month-old young SAMP8 mice.

## Results

### The pathophysiology of exercise, supplementary resveratrol intake, and combined of exercise and resveratrol has regulation at the 3- and 6-month-old SAMP8 mice

The aging liver appears a progressive decline of liver functions. Aging is characterized with morphological changes such as decrease in size and decreased hepatic blood flow. The most well-recognized risk for chronic diseases in physiological age. We used the SAMP8 mice liver to carry out by the aging test of the liver organ to age. The pathophysiology of liver cross-sections was detected by H&E, Masson’s trichrome, and TUNEL staining, treatment exercise, supplementary resveratrol intake, and combined of exercise and resveratrol by SAMP8 mice aging. Results showed adipocytes occurs in SAMP8 mice aging sections after exercise, supplementary resveratrol intake, and combination of exercise and resveratrol intake, the adipocytes become less. Especially, exercise training or resveratrol intake combination groups has recovered (Figure 1). Liver slicing is the quantification of fibrosis by H&E and Masson’s trichrome staining. In the 3- and 6-month-old SAMP8 mice liver by exercise, resveratrol, and the combination reduced adipocytes and collagens has significant differences (Figure 1A and 1B). Based on this finding, furthermore, apoptosis cells were determined by TUNEL assay. Exercise, supplementary resveratrol, and the combination facilitate the apoptosis cells reduced at the 3- and 6-month-old SAMP8 mice liver (Figure 2). In the 3- and 6-month-old SAMP8 mice liver, exercise, supplementary resveratrol, and the combination of exercise and resveratrol were reduced apoptosis cells (Figure 2). Especially, combined exercise and resveratrol have the best function to suppress apoptosis cells increases (p<0.001). Therefore, we found that combination of exercise and resveratrol can suppress aging-related liver diseases and enhance anti-aging.

**Figure 1.**
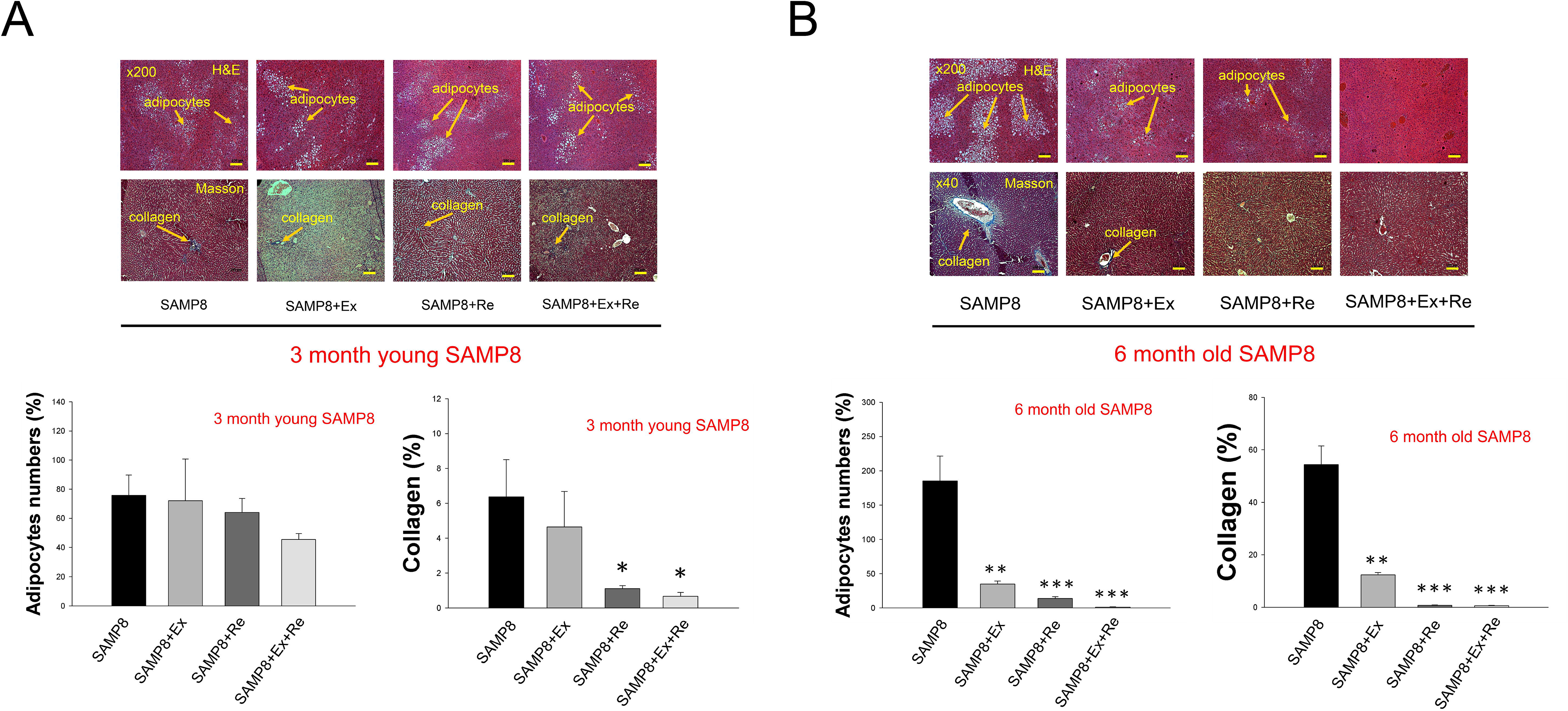
Histological pathophysiology of liver cross sections by exercise, resveratrol and the combination in the 3- and 6-month-old SAMP8 mice liver. Hematoxylin-eosin and Masson’s trichrome staining of liver cross sections by exercise, supplementary resveratrol intake and the combination (A) in the 3-month-old (young) and (B) 6-month-old (old) SAMP8 mice liver. Yellow arrow express adipocytes (above the lattice). Yellow arrow express collagen (below the lattice). Quantification of adipocytes numbers and collagen area densitometry analysis. All results are presented as mean ± SEM, \**p*<0.05, \*\**p<*0.01, \*\*\**p<*0.0001 significant difference compared with SAMP8 mice. Ex: exercise; Re: supplementary resveratrol intake; Ex+Re: combination of exercise and supplementary resveratrol.

**Figure 2.**
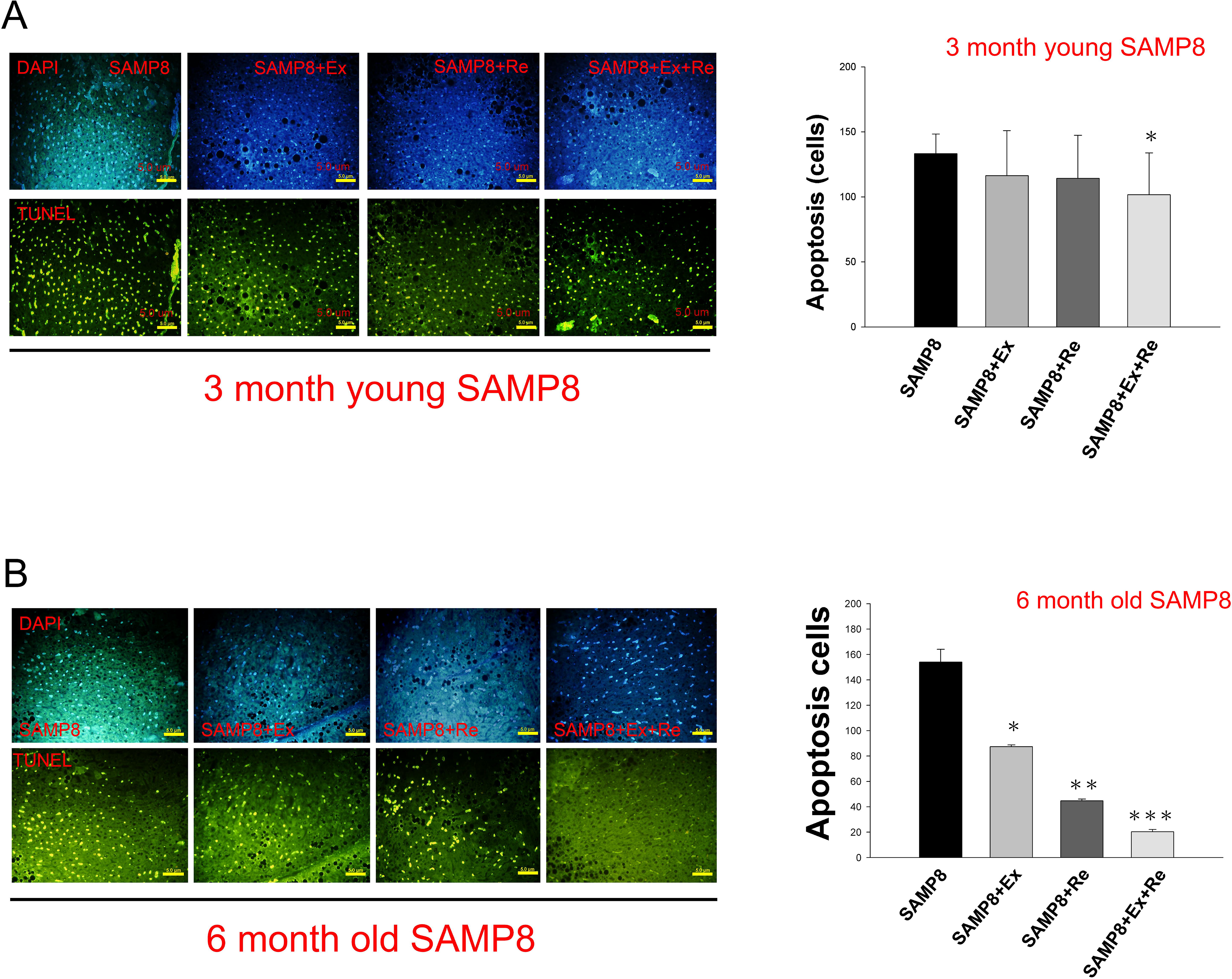
Apoptosis cell numbers of liver cross sections by exercise, supplementary resveratrol and the combination by TUNEL analysis in the 3- and 6-month-old SAMP8 mice liver. Cell apoptosis numbers in the (A). 3-month-old (young) and (B). 6-month-old SAMP8 mice liver (old) after exercise, supplementary resveratrol and the combination. Quantification of apoptosis cell numbers. All results are presented as mean ± SEM, \**p*<0.05, \*\**p<*0.01, \*\*\**p<*0.0001 significant difference compared with SAMP8 mice. Ex: exercise; Re: supplementary resveratrol intake; Ex+Re: combination of exercise and supplementary resveratrol.

### Exercise, supplementary resveratrol intake, and combined exercise and resveratrol facilitate PI3K/Akt signaling in 3- and 6-month-old SAMP8 mice liver

To identify senescence effect on hepatocytes survival signals PI3K/Akt ratios, in different degree SAMP8 mice liver. Protein expression levels of p-PI3K/PI3K and p-Akt/Akt ratios were detected in the 3- and 6-month-old SAMP8 mice liver by western blotting analysis (Figure 3). In the 3- and 6-month-old SAMP8 mice liver, we found p-PI3K/PI3K and p-Akt/Akt ratios have significant increases differences by the combination. In the 6-month-old SAMP8 mice liver has lower p-PI3K/PI3K ratio by exercise training compared with 3-month-old (p<0.05), but by supplementary resveratrol has lower p-Akt/Akt ratio (p<0.05). When at the 3-month-old SAMP8 mice compared with the 6-month-old SAMP8 mice, the combination of exercise and supplementary resveratrol has higher increases by 6-month-old SAMP8 mice liver (p<0.001) (Figure 3A). There may be because of mature genetic type of genetic senescence-accelerated. Exercise training and resveratrol intake have no significant difference at the 6-month-old SAMP8 mice. Resveratrol intake increased p-Akt/Akt ratio observed in the 3-month-old SAMP8 mice (p<0.05) (Figure 3B). In spite of resveratrol supplementary or exercise, there is no significant difference in the 6-month-old SAMP 8 mice liver. Therefore, combining with exercise and resveratrol supplement intake has good functions of p-Akt/Akt and p-PI3K/PI3K ratios in the 3- and 6-month-old SAMP8 mice liver (p<0.01).

**Figure 3.**
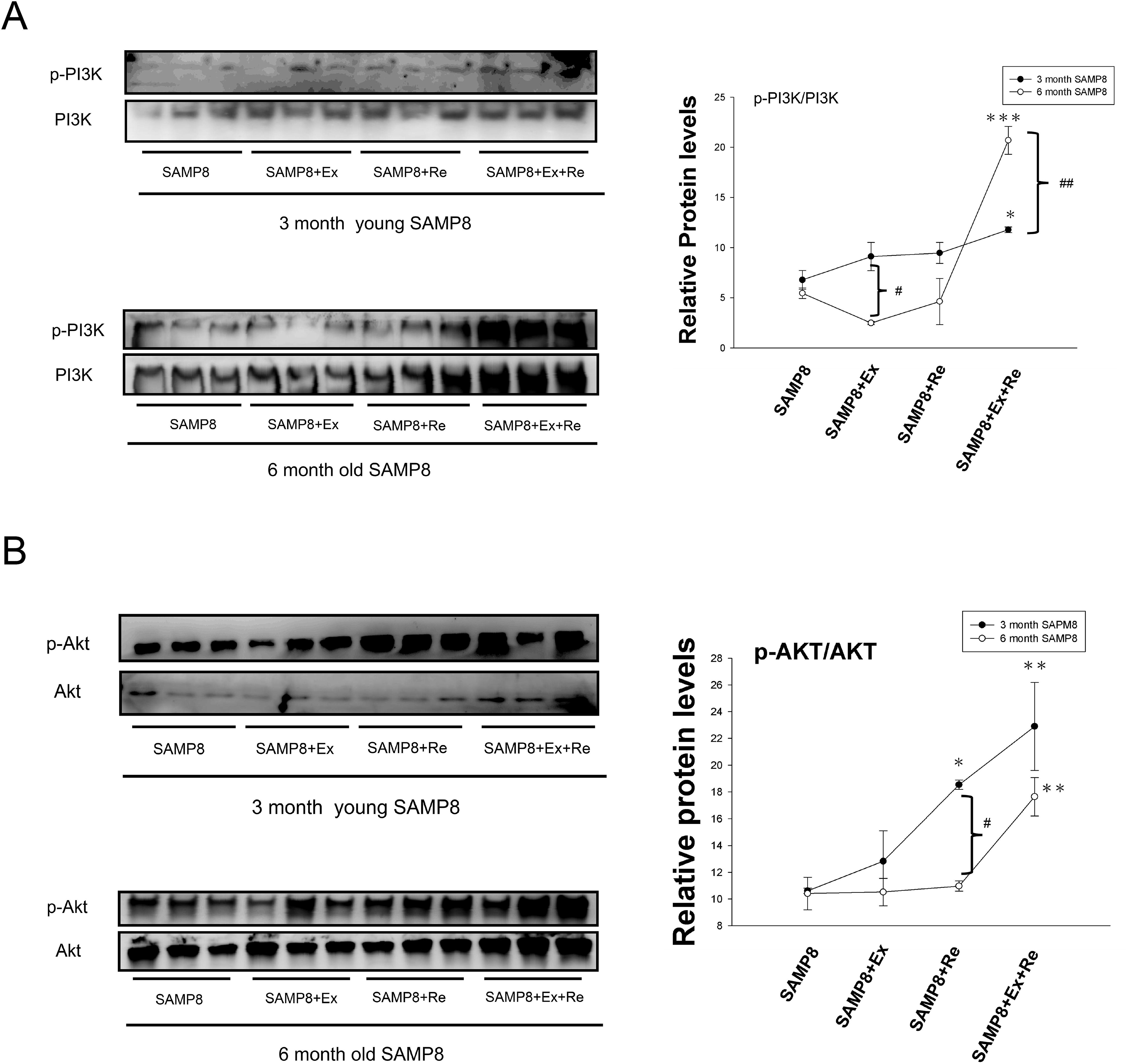
Regulation of cell survival signaling of p-PI3K/PI3K and p-AKT/AKT ratios increased by exercise, supplementary resveratrol intake and the combination in the 3- (young) and 6-month-old (old) SAMP8 mice liver. Protein expression levels of (A) p-PI3K/PI3K ratio and (B) p-AKT/AKT ratio after exercise, supplementary resveratrol and the combination in the 3- and 6-month-old SAMP8 mice liver by western blotting analysis. Quantification of densitometry analysis of protein expression levels. All results are presented as means ± SEM, \**p*<0.05, \*\**p<*0.01, \*\*\**p<*0.0001 significant difference compared with SAMP8 mice liver control. ^#^*p*<0.05, ^##^*p<*0.01 significant difference compared with the 3- and 6-month-old SAMP8 mice liver each other. Ex: exercise; Re: supplementary resveratrol intake; Ex+Re: combination of exercise and supplementary resveratrol.

### Increased mitochondrial capacity takes place during exercise, nutrition supplementary resveratrol, and combination of exercise and supplementary resveratrol in the age adaptations process

Mitochondrial related apoptosis during aging process is examined as follows. To detected mitochondrial cell death-related proteins, bad and cytochrome c, and anti-apoptosis protein, Bcl 2 and ERK1, in the 3- and 6-month-old SAMP8 mice liver by western blotting analysis (Figure 4 and 5). Results showed bad protein expression levels in the 3-month-old SAMP8 mice liver has no significantly different by all treatments, but in the mature degree of 6-month-old SAMP8 mice all treatments has significant decreases differences (Figure 4A). Nutrition resveratrol supplementary intake, exercise and combination of exercise and resveratrol were reduced cytochrome c in the 6-month-old SAMP8 mice liver (p<0.01) (Figure 4B). There are no significantly different by resveratrol supplementary intake and exercise in the 3-month-old SAMP8 mice liver. Once they are combination, we found had significant decreases (p<0.05). Compared with 3- and 6-month-old SAMP8 mice liver, cytochrome c has significantly different by exercise (p<0.01), nutrition supplementary resveratrol (p<0.05), and the combination (p<0.05). On the other hand, exercise induced Bcl-2 increased in the 3-month-old SAMP8 mice liver (p<0.01) (Figure 5A), but supplementary resveratrol and the combination did not induce Bcl-2 increases. However, the 6-month-old SAMP8 mice liver by exercise, supplementary resveratrol, and the combination has significant increases differences by Bcl-2 protein expression level increases (p<0.01). Compared with 3- and 6-month-old SAMP8 mice liver, mature 6-month-old has higher than 3-month-old by supplementary resveratrol and the combination (p<0.01). Interestingly, ERK1 has the same lower expression than Bcl 2 in the 3-month-old SAMP8 mice by exercise, resveratrol and their combination. However, in the 6-month-old SAMP8 mice liver by the combination has significant ERK1 protein expression increases effect (p<0.05). Exercise and resveratrol intake has no significant difference (Figure 5B).

**Figure 4.**
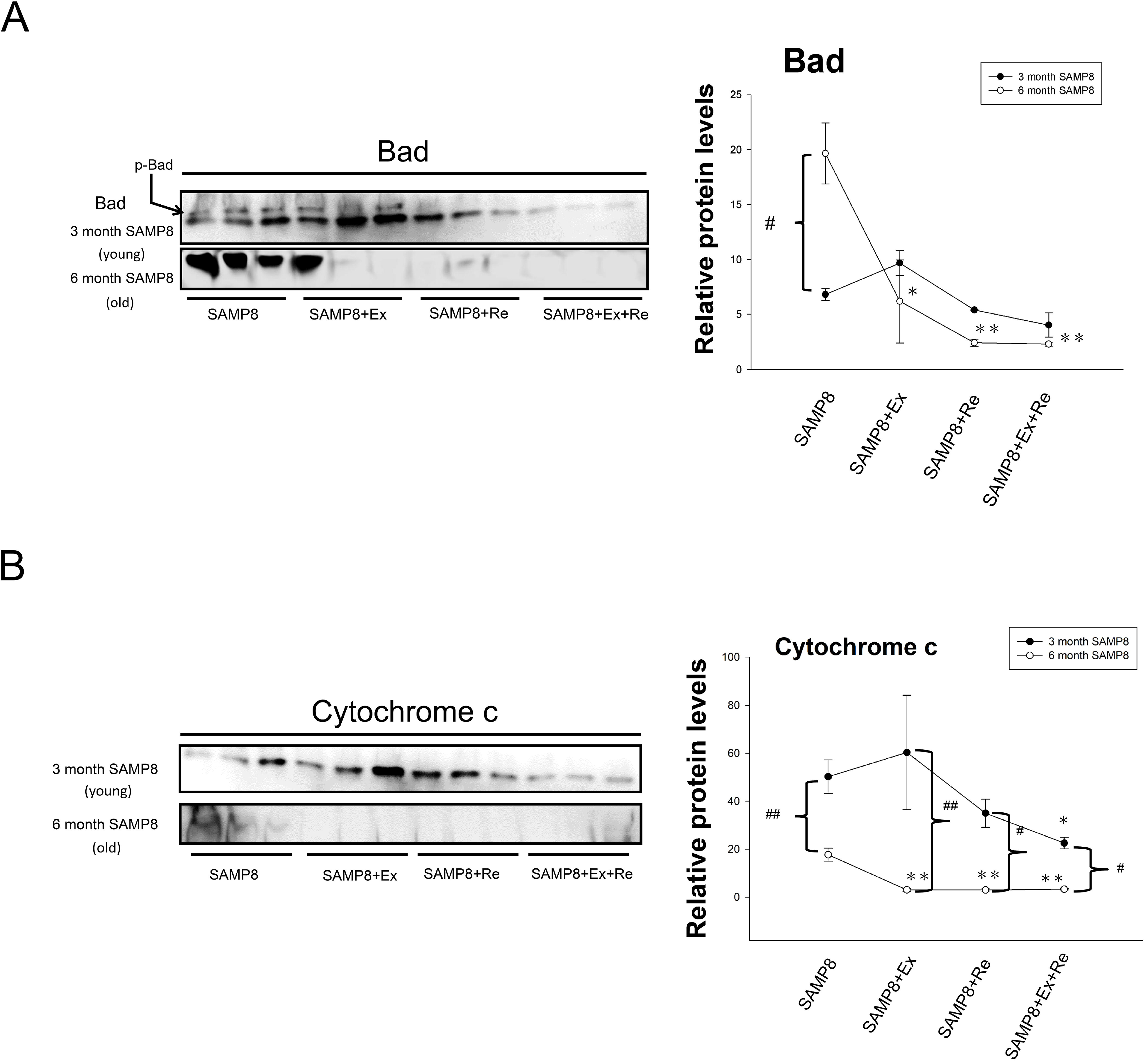
Regulation of mitochondrial apoptosis proteins, Bad and Cytochrome c, by exercise, supplementary resveratrol intake and the combination in the 3- (young) and 6-month-old (old) SAMP8 mice liver. (A). Protein expression of mitochondrial apoptosis, Bad, after exercise, resveratrol and the combination in the 3- and 6-month-old SAMP8 mice liver by western blotting analysis. (B). Protein expression of mitochondrial apoptosis, Cytochrome c, after exercise, supplementary resveratrol intake and the combination in the 3- and 6-month-old SAMP8 mice liver by western blotting analysis. All results are presented as means ± SEM, \**p<*0.05, \*\**p<*0.001 significant difference compared with SAMP8 mice. ^#^*p*<0.05, ^##^*p<*0.01 significant difference compared with the 3- and 6-month-old SAMP8 mice liver each other. Ex: exercise; Re: supplementary resveratrol intake; Ex+Re: combination of exercise and supplementary resveratrol.

**Figure 5.**
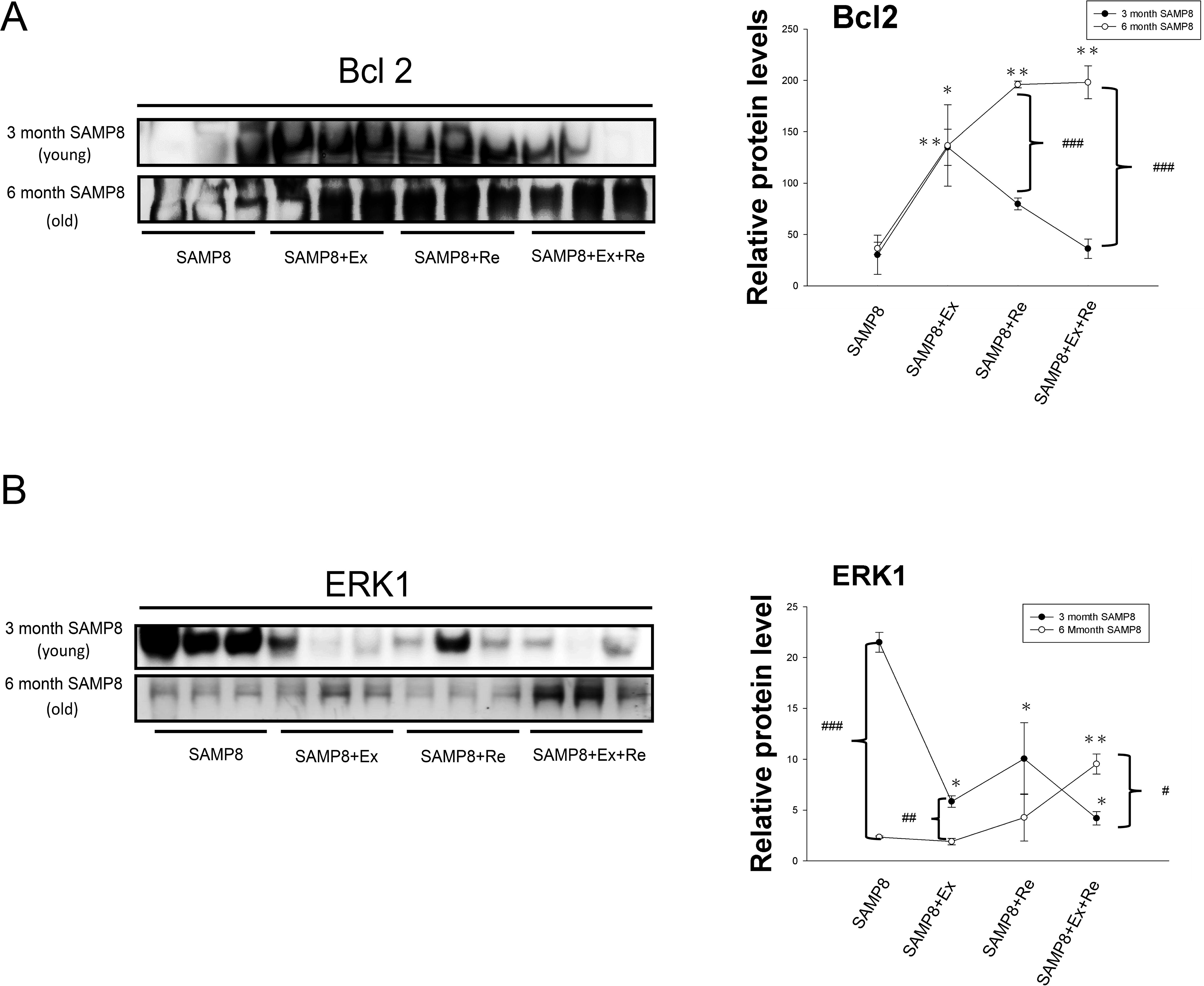
Regulation of cell proliferation signaling, ERK1, and mitochondrial anti-apoptosis, Bcl2, after exercise, supplementary resveratrol intake and the combination in the 3- (young) and 6-month-old (old) SAMP8 liver. (A). Protein expression of mitochondrial anti-apoptosis, Bcl2, after exercise, supplementary resveratrol intake and the combination in the 3- and 6-month-old SAMP8 mice liver. All results are presented as means ± SEM, \**p*<0.05, \*\**p<*0.01, significant difference compared with SAMP8 mice. (B). Cell proliferation protein expression levels of ERK1 by exercise, supplementary resveratrol intake and the combination in the 3- and 6-month-old SAMP8 mice liver by western blotting. Quantification of densitometry analysis of protein expression levels. All results are presented as means ± SEM, \**p*<0.05, \*\**p<*0.01, significant difference compared with SAMP8 mice. ^#^*p*<0.05, ^##^*p<*0.01, ^###^*p<*0.01significant difference compared with the 3- and 6-month-old SAMP8 mice liver each other. Ex: exercise; Re: supplementary resveratrol intake; Ex+Re: combination of exercise and supplementary resveratrol.

### Proliferation signaling pathway of IL-6/STAT3/MEK5/p-ERK5/ERK5 cascade in the 3- and 6-month-old SAMP8 mice liver

Inflammation factors, IL-6/STAT3/MEK5/p-ERK5/ERK5 cascade in the 3-month-old SAMP8 mice liver no significant difference by exercise, nutrition resveratrol intake, and combination of exercise training and nutrition resveratrol intake (Figure 6A). Nutrition resveratrol intake reduced IL-6 (p<0.05), but the combination of exercise and nutrition resveratrol intake has a little increase, but no significant difference compared with SAMP8 mice liver. The similar result was found at MEK5. Thus, exercise training and resveratrol intake could find significant reduced IL-6 protein expression (p<0.05). Their combination also did not find significant difference increases. Exercise, supplementary resveratrol intake, and the combination of exercise and resveratrol were decreases by STAT3 and p-ERK5/ERK5 ratio. STAT3 decreases have a significant difference by exercise and resveratrol (p<0.05). Their combination also did not find a significant difference decreases. The p-ERK5/ERK5 ratio has found significant difference decreases by combination and exercise training (p<0.05). But exercise did not find a significant difference. In the 6-month-old SAMP8 mice liver, we could find IL-6 and STAT3/MEK5/p-ERK5/ERK5 cascade decreased by exercise, nutrition resveratrol, and combination (Figure 6B). Remarkably, the combination has significantly reduced at IL-6 and STAT3/p-ERK5/ERK5 cascade.

**Figure 6.**
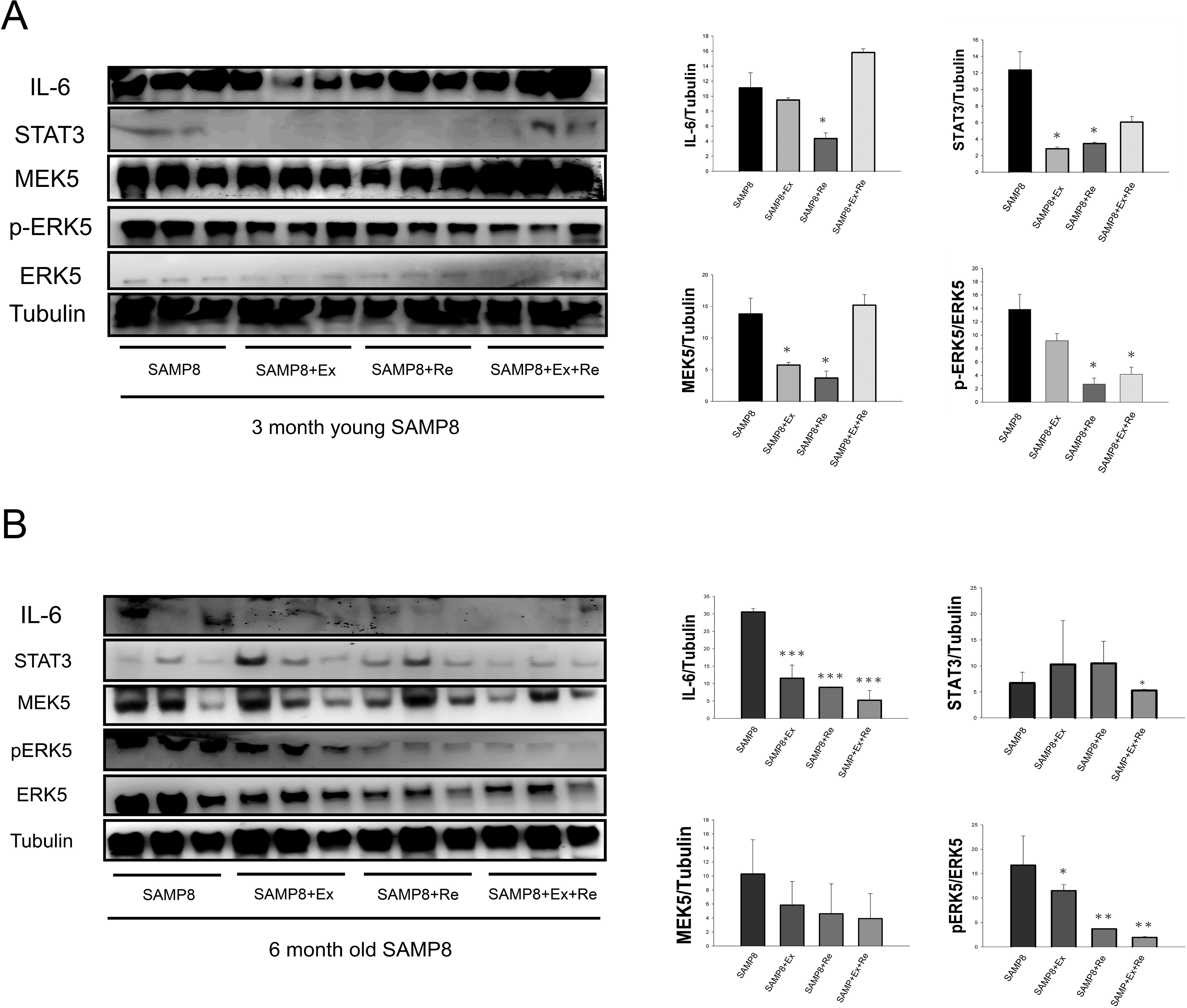
Regulation of inflammatory response system by western blot in aging liver after exercise, supplementary resveratrol intake and the combination. Inflammatory system and cell growth of IL-6/STAT3 and MEK5/p-ERK5/ERK5 protein expression levels after exercise, supplementary resveratrol intake and the combination by western blotting analysis in the (A). 3-month-old (young) and (B). 6-month-old (old) SAMP8 liver. Quantification of densitometry analysis of protein expression levels. All results are presented as means ± SEM, \**p*<0.05, \*\**p<*0.01, \*\*\**p*<0.0001 significant difference compared with SAMP8 mice. Ex: exercise; Re: supplementary resveratrol intake; Ex+Re: combination of exercise and supplementary resveratrol.

### Fibrosis signals of FGF2/MMP2/MMP9 by exercise, supplementary resveratrol intake, and combination in the 3- and 6-month-old SAMP8 mice liver

Downregulation of FGF2 in the 3- and 6-month-old SAMP8 mice liver, but upregulation of MMP2/MMP (Figure 7). Combination of exercise and resveratrol was no significant decreased by FGF2, but has a significant increase by MMP2 (p<0.05) in the 3-month-old SAMP8 mice liver (Figure 7A). However, FGF2 has significantly decreases by exercise, supplementary resveratrol, and combination (p<0.05) in the 6-month-old SAMP8 mice liver (Figure 7B). Exercise and resveratrol intake did not observed increased by MMP2/MMP9, but found at 3-month-old SAMP8 mice liver, respectively. Combination of exercise and resveratrol has a better FGF2/MMP2/MMP9 effect than exercise training or resveratrol intake.

**Figure 7.**
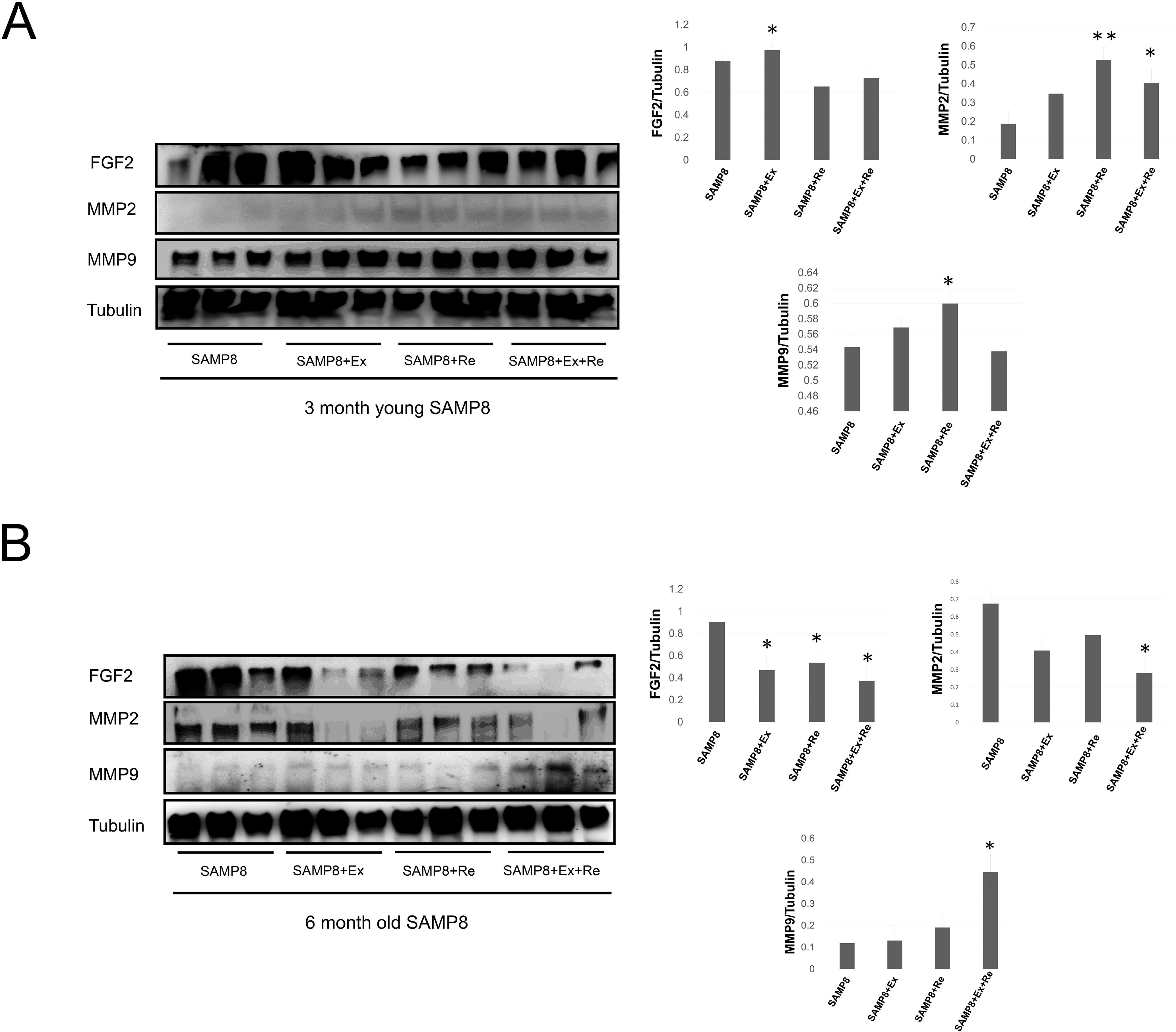
Western blotting of fibrosis proteins, FGF2 and MMP2/MMP9, in the 3- (young) and 6-month-old (old) SAMP8 mice liver. Fibrosis related proteins, FGF2 and MMP2/MMP9, in the (A). 3-month-old and (B). 6-month-old SAMP8 mice liver after exercise, supplementary resveratrol intake and the combination. Quantification of densitometry analysis of protein expression levels. All results are presented as means ± SEM, \**p*<0.05, \*\**p<*0.01, significant difference compared with SAMP8 mice. Ex: exercise; Re: supplementary resveratrol intake; Ex+Re: combination of exercise and supplementary resveratrol.

## Discussion

Life shortened occur as genetic alterations, such as DNA damage and cell suicide increase (Anversa et al., 1986a, 1990b). Aging liver is a gradually loses the ability maintained liver function due to liver injury. (Dolinsky and Dyck, 2014). The aging of the liver is not a disease, and the aging liver maintains its sound functions (Forman et al., 2013). To be fair, aging may be the process by which many factors participate together, and heredity and various environmental factor also play a decisive role (Halade et al., 2013; Roffe, 1998). However, aging liver is the reduction of liver functions because of adipocytes and collagen occurs (Figure 1). Due to unfavorable conditions like morbidities accompany it (Strait and Lakatta, 2012). Different stages of life or the pace of aging and performance of different, it can not be a single or simple model to be described (Olesen et al., 2014) With the increase in age, there are two cases of age-related physiological function decline, one for age-related physiologic deterioration (Olesen et al., 2013; Dolinsky et al., 2014), and age-associated disease with age, but two often co-exist (Kwak, 2013). Aging liver is one of age-related physiologic deterioration (Yu et al., 2013). Regular exercise training maybe improves, and resveratrol intake may cause changes. Exercise, supplementary resveratrol intake and the combination facilitate the interaction of cell death and its life span in the 3- and 6-month-old SAMP8 mice liver. Age-related liver disease ameliorate apoptosis by exercise, resveratrol and combination. We could observed reversed apoptosis, collagen and adipocytes effects on by exercise, resveratrol and combination of excise and resveratrol (Figure 1 and 2). Accelerated different aging grade induced different degree cell life and death interaction lead to unbalance of p-PI3K/PI3K and p-Akt/Akt ratios, and Bad-Bcl2-Cytochrome c (Figure 3, 4 and 5). Exercise, resveratrol and the combination resulted in different regulation of PI3K-Akt-ERK1-Bcl-2 and Bad-Bcl-2-Cytochrome c (Vetterli and Maechler, 2011; Hou et al., 2016). Physical exercise is for maintaining physical fitness including healthy (Tung et al., 2015; Bezzerides and Rosenzweig, 2011), building and maintaining healthy age liver (Tung et al., 2014); promoting physiological well-being; reducing aging risks (Tung et al., 2015); and strengthening the inflammatory response (Momchilova et al., 2014; Hamer et al., 2010). The immune system of IL-6 and STAT3/MEK5/ERK5 was stretching improve in the 6-month-old SAMP8 mice liver (Figure 6). Exercise, supplementary resveratrol and the combination in the 3-month-old SAMP8 liver has a little anti-fibrosis effects function (Figure 7A). However, combination exercise and resveratrol has excellent function by FGF2/MMP2/MMP9 in the 6-month-old SAMP8 mice liver (Figure 6B).

## Conclusions

Resveratrol supplements can slow the aging process and promote longer cell life in the body (Boström et al., 2010; Bernardo et al., 2010; Chai, et al., 2017; Mukherjee et al., 2009; Safdar et al., 2011). Genetic senescence-accelerated aging induces age-related liver disease by exercise training, resveratrol intake and the combination ameliorate aging liver apoptosis (Momchilova et al., 2014; Yang and Lim, 2014; Das et al., 2016). Exercise, resveratrol and the combination ameliorated different degree aging for 6-month-old is better than 3-month-old SAMP8 mice liver. Therefore, the aging liver can be regarded as a latent liver disease factor in countries or communities where malnutrition and infections are the prevalent long-term 6-month-old SAMP8 mice liver.

## Abreviations

H&E: hematoxylin-eosin staining
TBS: Tris-buffered saline
BSA: bovine serum albumin
SAMP: senescence-accelerated mice

## Acknowledgments

This work was supported in part by grants from the national science council of Taiwan.

## Funding

No available.

## Competing interests

No competing interest exists.

